# Exploring the Comedone Switch through Single-Cell Resolution of Human Acne Lesions

**DOI:** 10.64898/2026.07.09.737640

**Authors:** Tolga Düz, Tobias Rolka, Daniel Torocsik, Hendrik Reuter, Benjamin Al, Stefan Gallinat, Jan Baumbach, Nicholas Holzscheck

**Author notes:** Corresponding author: Tolga Düz, Beiersdorfstraße 1-9, 20245 Hamburg, Germany. Emails and ORCIDs: Tolga Düz –; Tobias Rolka –; Daniel Torocsik –; Hendrik Reuter –; Benjamin Al –; Stefan Gallinat –; Jan Baumbach –; Nicholas Holzscheck –.

## Abstract

Acne vulgaris is one of the most prevalent inflammatory skin diseases worldwide, yet the molecular events initiating comedogenesis remain poorly understood. The comedone switch hypothesis proposes that acne originates from an imbalance in lineage commitment within the junctional zone of the pilosebaceous unit, promoting infundibular differentiation at the expense of sebaceous gland maintenance. However, direct evidence from human acne tissue at single-cell resolution has been lacking. Here, we integrated single-cell transcriptomic datasets from healthy skin, nonlesional skin of acne patients, and lesional acne tissue to investigate early transcriptional changes associated with comedogenesis. We identified a previously uncharacterized cell population in nonlesional skin with transcriptomic features suggestive of a microcomedone-like state and mapped this population across independent datasets to characterize its relationship to comedonal keratinocytes. Comedonal remodeling was characterized by enhanced keratinization and inflammatory programs. Quantitative analyses supported a shift from sebaceous toward infundibular cell fate, providing data-driven support for the comedone switch hypothesis in human acne. Beyond the pilosebaceous unit, we identified broader epithelial alterations, including loss of POSTN and ERRFI1 expression in basal interfollicular epidermal keratinocytes. Together, these findings provide a cell-resolved view of transcriptional changes associated with human comedogenesis and highlight candidate pathways that may contribute to lineage imbalance within the upper hair follicle.

## INTRODUCTION

Acne vulgaris is one of the most common skin diseases, affecting approximately 650 million individuals worldwide (Bhate and Williams, 2013; Vos et al., 2012). Despite its high prevalence and significant clinical, psychological, and economic burden (Halvorsen et al., 2011; Hazarika and Archana, 2016; Xu et al., 2021; Zaraa et al., 2013), the pathogenesis of acne remains incompletely understood.

Acne originates within the pilosebaceous unit (PSU), a skin appendage comprising the hair follicle and the sebaceous gland (SG) (Tuchayi et al., 2015). The upper PSU opens at the infundibulum, which transitions into the isthmus. The junctional zone denotes the region where the SG connects to the isthmus via the sebaceous duct (SD) (Figure 1a) (Schneider et al., 2009). The SG is a multilobular holocrine gland with basal proliferative cells at the periphery and differentiating sebocytes at the center. As they mature, sebocytes accumulate lipids and rupture, releasing sebum into the upper PSU (Fischer et al., 2017; Schneider and Paus, 2010).

**Figure 1:**
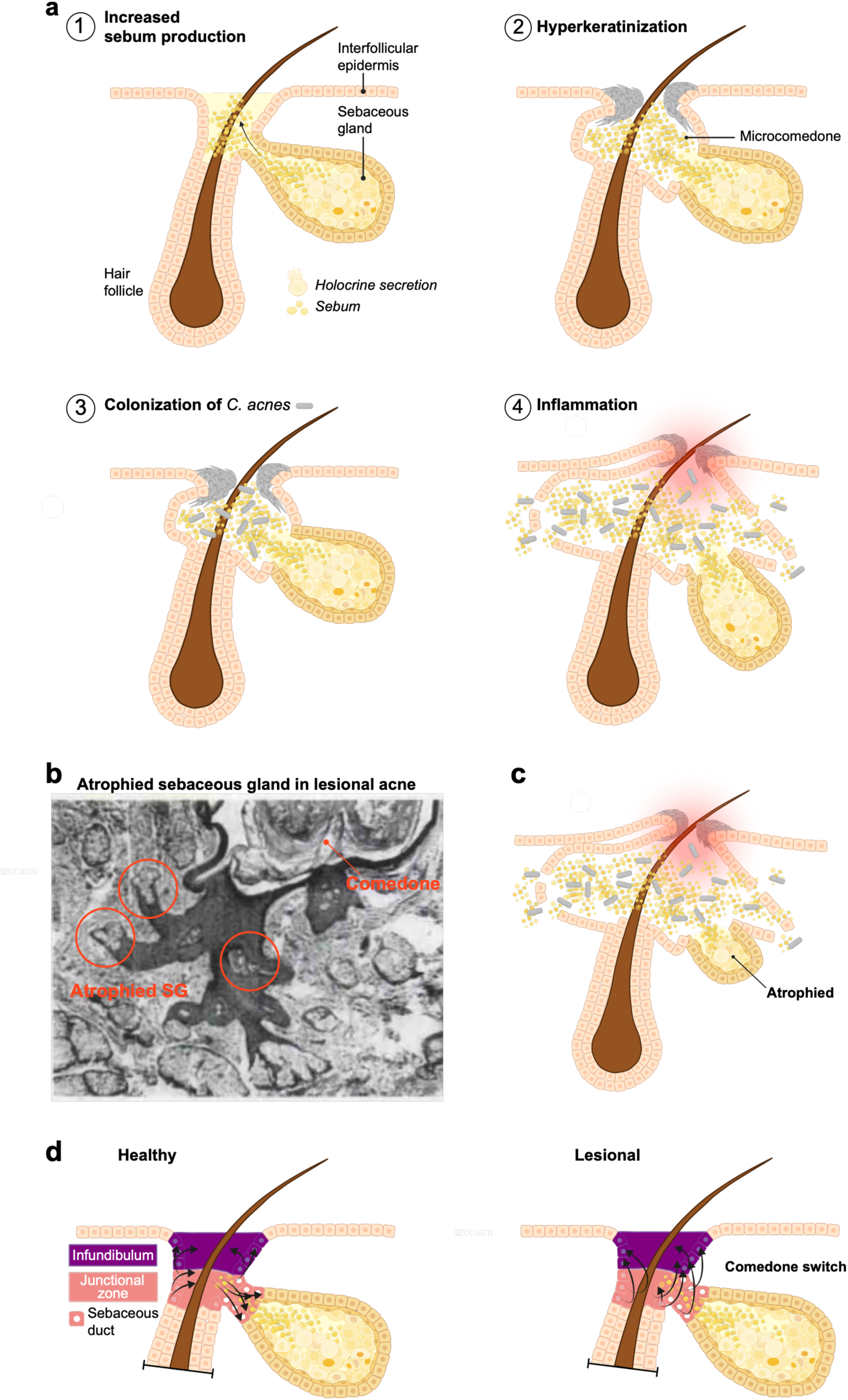
Canonical hallmarks of acne pathogenesis and the comedone switch hypothesis. **(a)** Schematic overview of the human PSU, consisting of the hair follicle and SG. Four canonical factors have been implicated in acne pathogenesis: increased sebum production, follicular hyperkeratinization, colonization by *Cutibacterium acnes*, and inflammation. **(b)** Histological image of acne-affected skin demonstrating sebaceous gland atrophy associated with comedone formation. **(c)** Schematic illustration of an atrophied sebaceous gland within an inflamed comedone. **(d)** Schematic representation of the comedone switch hypothesis. Under homeostatic conditions, bipotent progenitor cells within the junctional zone maintain a balance between sebaceous and infundibular differentiation. In acne, lineage commitment is proposed to shift toward infundibular hyperproliferation and differentiation at the expense of sebaceous gland maintenance, resulting in follicular occlusion, comedone formation, and progressive sebaceous gland atrophy.

Understanding this anatomy is essential, as several pathogenic factors are localized within these compartments. Traditionally, four major factors have been implicated (Figure 1a) (Vasam et al., 2023):

i. increased and altered sebum production under androgenic and hormonal control;
ii. hyperkeratinization of the follicular infundibulum, resulting in follicular occlusion that prevents sebum release and promotes comedone formation;
iii. colonization of the follicle by *Cutibacterium acnes*, and
iv. the release of inflammatory mediators.

While the four major factors are well established, their causal relationships and sequence of events remain unclear (Saurat, 2015). Furthermore, this conventional model offers limited explanatory power for several features of acne pathogenesis, such as why lesions develop in only a subset of PSU. This spatial selectivity indicates that lesion formation likely requires additional local triggers (Saurat, 2015; van Steensel and Goh, 2021).

Another inconsistency with the concept of globally increased and altered sebum production as the primary factor is the frequent atrophy of SGs in lesional acne skin, already described in histological studies as early as the 1970s (Figure 1b and c) (Kligman, 1974; Plewig, 1974). This atrophy seemingly contradicts the notion that increased sebum production is the primary driver of comedogenesis and remains difficult to reconcile with the four canonical factors (Clayton et al., 2019; van Steensel, 2025; van Steensel and Goh, 2021), hinting at a more complex pathogenesis than assumed by the classical disease model.

This is further supported by clinical observations in chloracne, a dioxin-induced condition where comedone-like cysts form in the complete absence of SGs (Saurat et al., 2012). These findings suggest that a differentiation defect in the upper hair follicle epithelium, rather than sebaceous hyperactivity, may represent the primary pathogenic event in acne.

Based on these observations and supported by evidence that cells in the JZ possess bipotent potential to adopt either an infundibular or sebaceous fate (Jensen et al., 2009), the comedone switch hypothesis was proposed (Saurat, 2015). This model posits that acne results from a shift in lineage commitment within a common progenitor population in the JZ, favoring infundibular hyperproliferation and differentiation at the expense of SG maintenance (Figure 1d). As a result, the follicle becomes clogged, and the basal cell layer of the SG is no longer replenished, ultimately leading to its disappearance (Clayton et al., 2019; Saurat, 2015; van Steensel, 2025; van Steensel and Goh, 2021).

This hypothesis provides a plausible explanation for SG atrophy and incorporates a local component that makes certain PSUs more susceptible, potentially influenced by genetic predispositions, thereby explaining the spatial restriction of lesions. However it has remained without much concrete evidence until recently.

To investigate whether Wnt signaling functions as a molecular switch governing lineage decision of bipotent JZ stem cells, Shang et al. (2022) used the Lrig1-CreERT2 mouse line to modulate Wnt activity in vivo. Gain-and loss-of-function experiments revealed that persistent Wnt activation, induced by Apc deletion, leads to cyst formation and SG atrophy. In contrast, Wnt inhibition through β-catenin deletion promoted balanced differentiation while preserving tissue structure.

Another study using a sebaceous organoid model demonstrated that the transcription factor GATA6 regulates upper hair follicle homeostasis by downregulating infundibular fate in the JZ through TGFβ activation (Oules et al., 2020). Its reduced expression in acne-affected skin suggests a potential role in acne pathogenesis by promoting the comedone switch.

However, acne is a human-specific disease, and no animal or in vitro model completely captures its complex pathophysiology. Moreover, the referenced organoid study relied on single-cell RNA sequencing (scRNA-seq) of healthy human skin and compared these data to bulk transcriptomic profiles from comedonal acne lesions. This cross-platform approach introduces batch effects and lacks cell-type-specific resolution in diseased tissue. As a result, the analysis cannot capture the cellular heterogeneity or identify acne-associated changes within specific cell populations.

A comprehensive, cell-resolved transcriptomic comparison between healthy, nonlesional, and acne-affected human skin represents one of the most powerful current approaches to elucidate the mechanisms underlying acne pathogenesis. Although several reviews have summarized key aspects of acne development (Lai et al., 2025; van Steensel, 2025; Vasam et al., 2023; Zouboulis, 2020), our understanding of human comedogenesis from microcomedone formation to its progression into inflamed lesions remains incomplete. Visible lesions ultimately emerge from the microcomedone (Fontao et al., 2020; Saurat, 2015), and a deeper understanding of its origin and development could therefore provide important insights into acne pathogenesis.

In this study, we leveraged scRNA-seq data from the nonlesional skin of acne patients identifying a previously uncharacterized cluster with transcriptomic features consistent with the microcomedone. Based on its gene expression signature, we mapped this putative microcomedone-like transcriptional state onto two independent lesional acne scRNA-seq datasets, enabling a molecular reconstruction of the comedone architecture at single-cell resolution. This approach allowed us to resolve the PSU in both nonlesional and acne-affected human skin, including the identification of sebocytes.

Furthermore, we developed a biologically interpretable model that captures the molecular determinants of the comedone switch and provides quantitative support for this concept in human acne lesions. By resolving PSU and IFE cell types under different conditions, we uncovered distinct gene expression profiles likely associated with comedogenesis and identified candidate upstream signaling pathways that may drive this process. Notably, several of these pathways are supported by SNPs identified in genome-wide association studies (GWAS).

Finally, our single-cell-based characterization of the putative comedone allowed us to critically evaluate previously proposed comedone markers from the literature. Our findings suggest that deciphering cell fate decisions within the PSU, particularly within the JZ, may open new therapeutic avenues for acne and other disorders of SG dysfunction.

## 2 RESULTS

### Fine-grained analysis of the keratinocyte compartment reveals a putative microcomedone-like transcriptional state in nonlesional acne skin

To investigate cellular changes underlying acne pathogenesis, we reanalyzed the public scRNA-seq dataset GSE175817 (Do et al., 2022), which includes both lesional and adjacent nonlesional back skin from six acne patients. Analysis of nonlesional skin provides a unique opportunity to identify early transcriptional changes that may precede the clinical onset of acne lesions.

We subclustered the keratinocyte compartment from nonlesional skin cells, to achieve higher resolution of transcriptionally distinct subpopulations. This approach enabled the delineation of functionally distinct cell states within the PSU and interfollicular epidermis (Do et al., 2022). The annotation of these populations was supported by expression patterns consistent with previous reports (Figure 2b).

**Figure 2:**
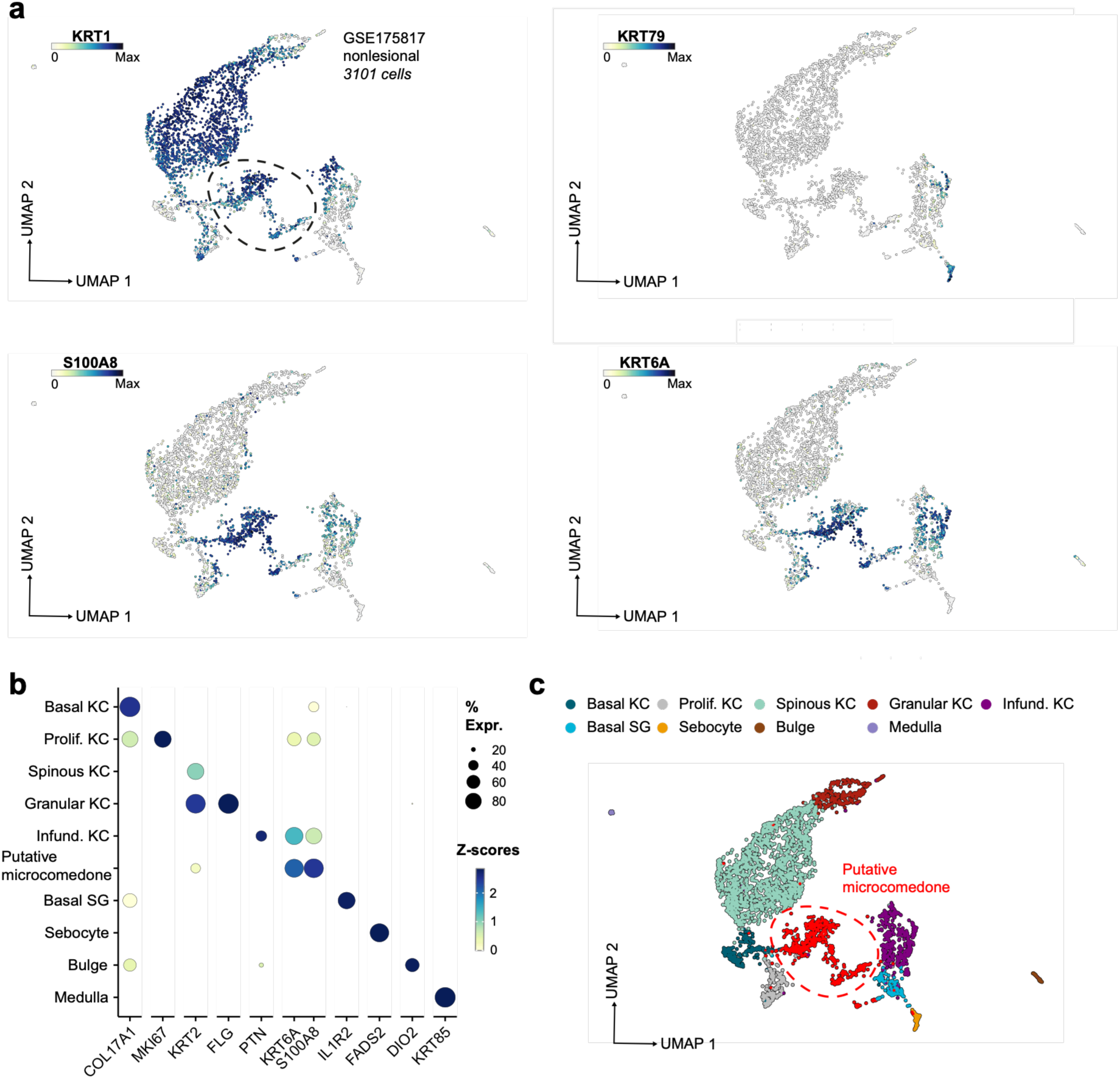
Identification of a putative microcomedone population in nonlesional acne skin. **(a)** UMAP representation of keratinocyte subclusters derived from single-cell RNA sequencing data of nonlesional skin from acne patients (GSE175817). **(b)** Dot plot showing the expression of selected marker genes used for cell type annotation. Expression patterns were consistent with previously reported markers of epidermal and PSU cell populations. **(c)** A previously unrecognized cluster located between infundibular and epidermal keratinocytes is highlighted (dashed circle). This cluster exhibited features of suprabasal differentiated keratinocytes while retaining partial transcriptional similarity to the infundibulum. Compared with healthy infundibular keratinocytes, these cells displayed elevated expression of inflammatory markers. Because this population was detected exclusively in nonlesional acne skin and was absent from healthy reference datasets, we hypothesize that it represents the microcomedone, the earliest precursor lesion in acne pathogenesis.

Notably, we identified a distinct cluster located between the infundibulum and epidermal keratinocyte clusters (indicated by a dashed circle in Figure 2a). This cluster expressed terminal differentiation markers, including KRT1 and IVL, consistent with a suprabasal keratinocyte identity, yet also displayed partial infundibular characteristics (Figure 2a). For example, PTN and KRT79, markers enriched in the infundibulum and junctional zone in healthy skin scRNA-seq datasets (Düz et al., 2026a; Düz et al., 2026b) , were not expressed in this population. Conversely, S100A8 and KRT6A, also enriched in the same regions in healthy skin, were expressed at higher levels compared to infundibular keratinocytes. This pattern suggests that the cluster does not represent the normal infundibular population, but rather keratinocytes undergoing aberrant differentiation or exhibiting an inflammatory activation state near the infundibulum. Importantly, this cluster was exclusively detected in nonlesional skin and absent from the healthy Human Skin Cell Atlas (HSCA) (Supplementary Figure S1a) (Düz et al., 2026a), showing high specificity to three patients (Supplementary Figure S1b).

Since acne primarily affects the PSU and the sampled nonlesional skin appears macroscopically healthy, we hypothesize that this cluster represents the **microcomedone** (Figure 2c), a macroscopically invisible precursor lesion. As an early detectable molecular alteration preceding visible comedone formation, this cluster is considered to mark an initial pathogenic event in acne development and may serve as a critical entry point for studying disease initiation (Saurat, 2015).

### Integration with lesional skin supports microcomedone identity through cross-dataset modeling

To support the hypothesis that the identified cell state in nonlesional skin corresponds to the microcomedone, we integrated nonlesional and matched lesional datasets from the same individuals (Figure 3a) (Do et al., 2022). In the resulting embedding, we observed a prominent lesional-specific cluster characterized by elevated expression of inflammatory and keratinization-associated genes within the upper hair follicle. Notably, the putative microcomedone-like transcriptional state from nonlesional skin localized within this lesional-specific region of the integrated embedding (Supplementary Figure S2). This positional continuity is consistent with the interpretation that the microcomedone represents an early, inflammatory precursor state that may give rise to the inflamed comedone in acne.

**Figure 3:**
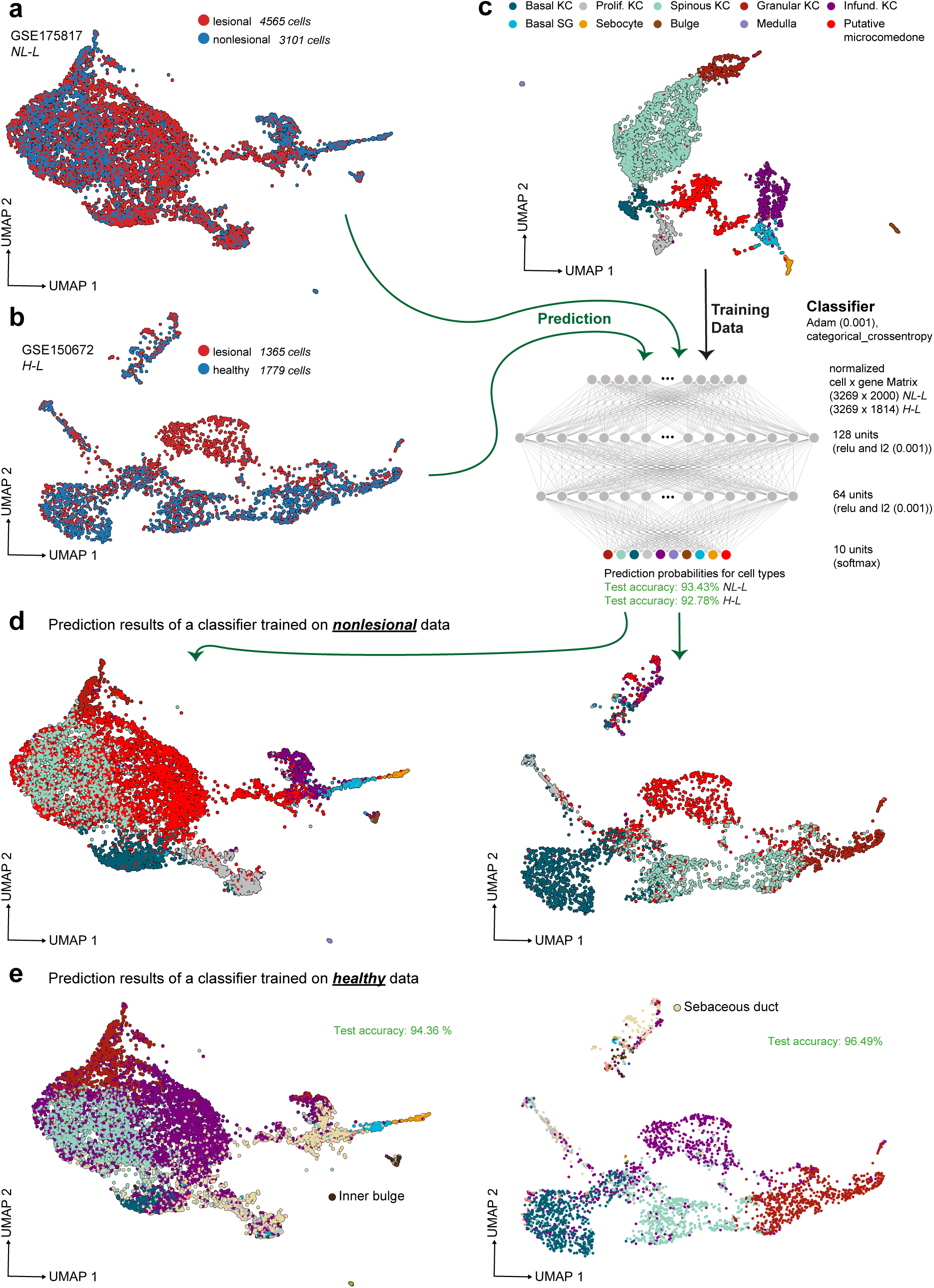
Cross-dataset validation of the putative microcomedone and reconstruction of acne-associated cell states. **(a)** Integrated embedding of nonlesional and matched lesional acne skin from GSE175817. The putative microcomedone-like transcriptional state identified in nonlesional skin localizes within a lesional-specific region of the embedding, suggesting transcriptional continuity between the two states. **(b)** Integrated embedding of healthy and lesional acne skin from an independent dataset. A distinct cluster composed exclusively of lesional cells was identified and characterized by elevated expression of inflammatory and keratinization-associated genes. **(c)** Schematic overview of the classification framework. A neural network classifier was trained on annotated nonlesional keratinocyte subclusters, including the putative microcomedone-like transcriptional state, and subsequently applied to integrated acne datasets. **(d)** Predicted cell state identities following classifier application. Lesional acne-associated clusters were predominantly assigned to the putative microcomedone state, supporting their transcriptional similarity. **(e)** Prediction of cellular origin using a classifier trained exclusively on healthy skin populations. Microcomedone-like cells were predominantly assigned an infundibular identity, suggesting that they arise from infundibular keratinocytes undergoing aberrant differentiation. Abbreviations: H-L, healthy vs. lesional; KC, keratinocyte; NL-L, nonlesional vs. lesional.

To further examine this phenomenon in an independent dataset comparing healthy and lesional acne skin (Hughes et al., 2020), we applied the same subclustering approach. In the integrated embedding, we observed a distinct cluster composed exclusively of cells from lesional samples, with no contribution from healthy skin (Figure 3b). This lesion-specific cluster exhibits elevated expression of inflammatory and keratinization-associated genes and may represent a later-stage, inflamed comedone, consistent with the absence of microcomedone-related cell states in healthy individuals.

To test whether the lesional clusters share transcriptional similarity with the putative microcomedone-like transcriptional state in a fully data-driven and unbiased manner, we trained two neural network classifiers. The first model was trained on annotated subclusters from nonlesional skin, including the previously identified putative microcomedone cluster (Figure 3c). Our rationale was that if this state represents a precursor to the comedone, lesional-specific clusters would be expected to show the highest transcriptional similarity to it. When applied to the integrated datasets, the classifier predominantly assigned the lesional clusters to the microcomedone-like state (Figure 3d).

To infer the cellular origin of comedonal cells, we trained a second model using only healthy skin samples, which represent unperturbed lineage programs. The classifier consistently predicted an infundibular identity for the microcomedone-like cells (Figure 3e), suggesting that they likely arise from infundibular keratinocytes undergoing aberrant differentiation. This distinction between transcriptional similarity and inferred lineage origin is consistent with our interpretation of the microcomedone as a pathological state emerging from the infundibular compartment.

Furthermore, both datasets were projected onto the core healthy HSCA using transfer learning (Düz et al., 2026a). This atlas comprises scRNA-seq data from approximately 300,000 cells covering a range of skin regions and encompassing diverse ages, sexes, and ethnicities. Transfer learning enabled automated cell-type annotation and provided an uncertainty metric, where high uncertainty indicates that a given cell state is not represented in the healthy reference atlas. Notably, putative comedonal keratinocytes exhibited the highest uncertainty scores across all keratinocyte populations, consistent with the absence of a corresponding cell state in healthy skin (Figure 4a and Supplementary Figure S3). In contrast, established epidermal, SG, and hair follicle cell populations showed substantially lower uncertainty scores, supporting robust mapping to healthy reference cell types. The final cell-type annotations are shown in Figure 4b, with corresponding marker gene expressions in Figure 4c.

**Figure 4.**
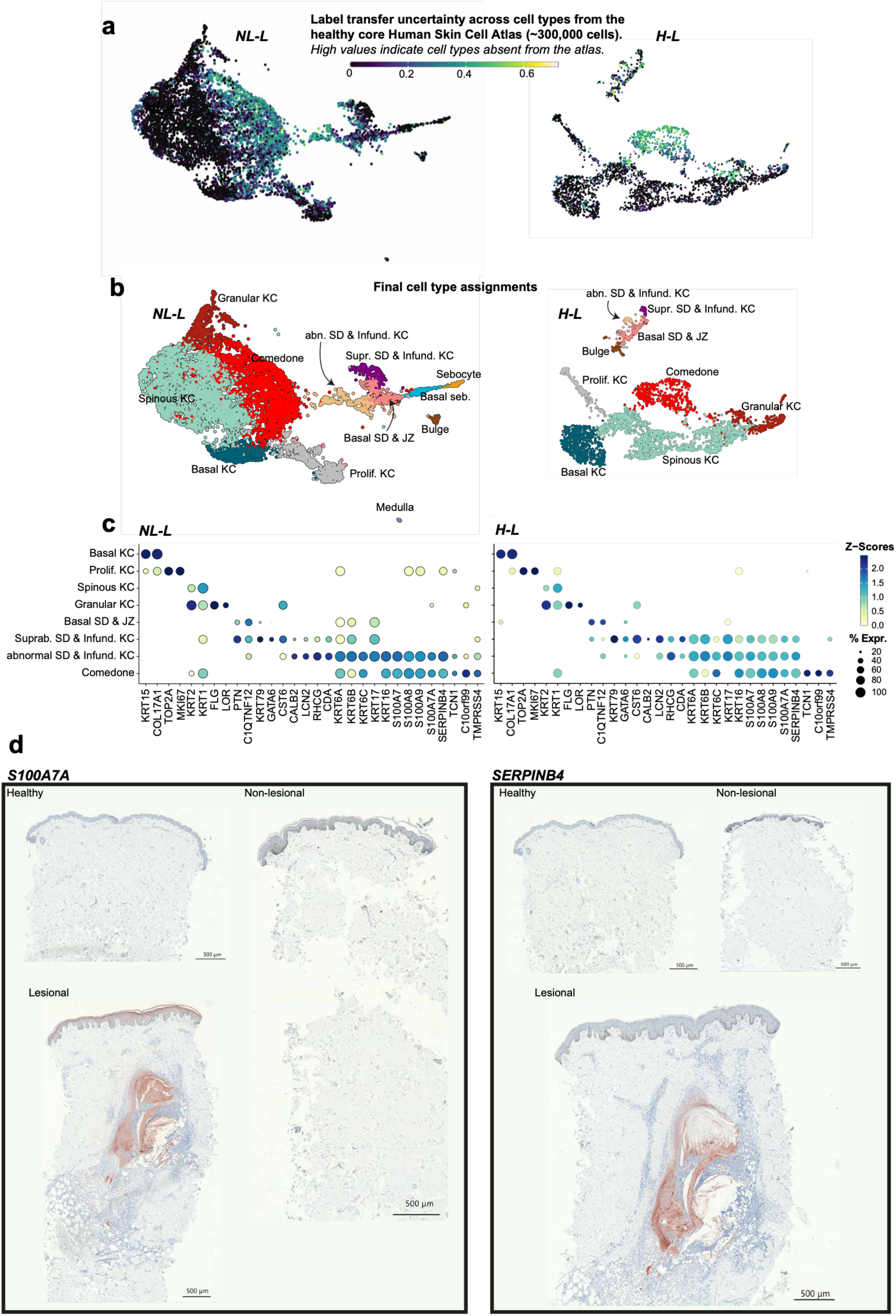
Transfer learning-based annotation of acne-associated cell states and validation of the comedone gene signature. **(a)** Projection of integrated acne datasets onto the healthy Human Skin Cell Atlas using transfer learning. Prediction confidence reflects the similarity of queried cells to healthy reference populations, with low confidence indicating cell states that are poorly represented or absent from healthy skin. **(b)** Final cell type annotations. **(c)** Expression of selected marker genes supporting the cell type annotations. **(d)** Immunohistochemical validation of the comedone-associated gene signature. Representative staining for S100A7A and SERPINB4 in healthy, nonlesional, and lesional acne skin. Strong expression was observed specifically within comedonal structures, supporting the association of these markers with the comedone cell state identified by single-cell RNA sequencing. Abbreviations: H-L, healthy vs. lesional; JZ, junctional zone; KC, keratinocyte; NL-L, nonlesional vs. lesional; SD, sebaceous duct.

To validate the molecular identity of comedonal keratinocytes in situ, we performed immunostaining (Figure 4d). As a first marker, we selected S100A7A. In healthy skin, S100A7A expression was not detected and remained absent in nonlesional skin. In contrast, acne lesions exhibited strong S100A7A staining. Notably, expression was observed not only in keratinocytes within the comedone but also in the overlying epidermis, indicating that S100A7A induction is not restricted to the comedonal structure. A similar expression pattern was observed for SERPINB4, which was absent in healthy and nonlesional skin but strongly expressed in acne lesions.

### Exploring the comedone switch hypothesis through lineage modeling of keratinocyte and sebocyte differentiation

Next, we investigated the cellular basis of comedogenesis by comparing lesional and nonlesional skin using our integrated single-cell datasets. Quantification of cell numbers revealed a marked depletion of SG cells in lesional skin (43 versus 152 cells), accompanied by a reduction in infundibular keratinocytes (158 versus 295 cells) and a substantial expansion of comedonal keratinocytes (1,785 versus 496 cells) (Supplementary Figure S4). These shifts were readily apparent in the integrated embedding (Figure 5a and b). This pattern is consistent with the comedone switch hypothesis (Saurat, 2015), potentially suggesting a shift in lineage commitment from sebocytes toward aberrantly differentiating infundibular keratinocytes, leading to SG atrophy and expansion of the keratinized comedonal compartment. Notably, the comedonal keratinocytes detected in nonlesional skin corresponded to the putative microcomedone-like transcriptional state identified in our previous analyses (Supplementary Figure S2).

**Figure 5.**
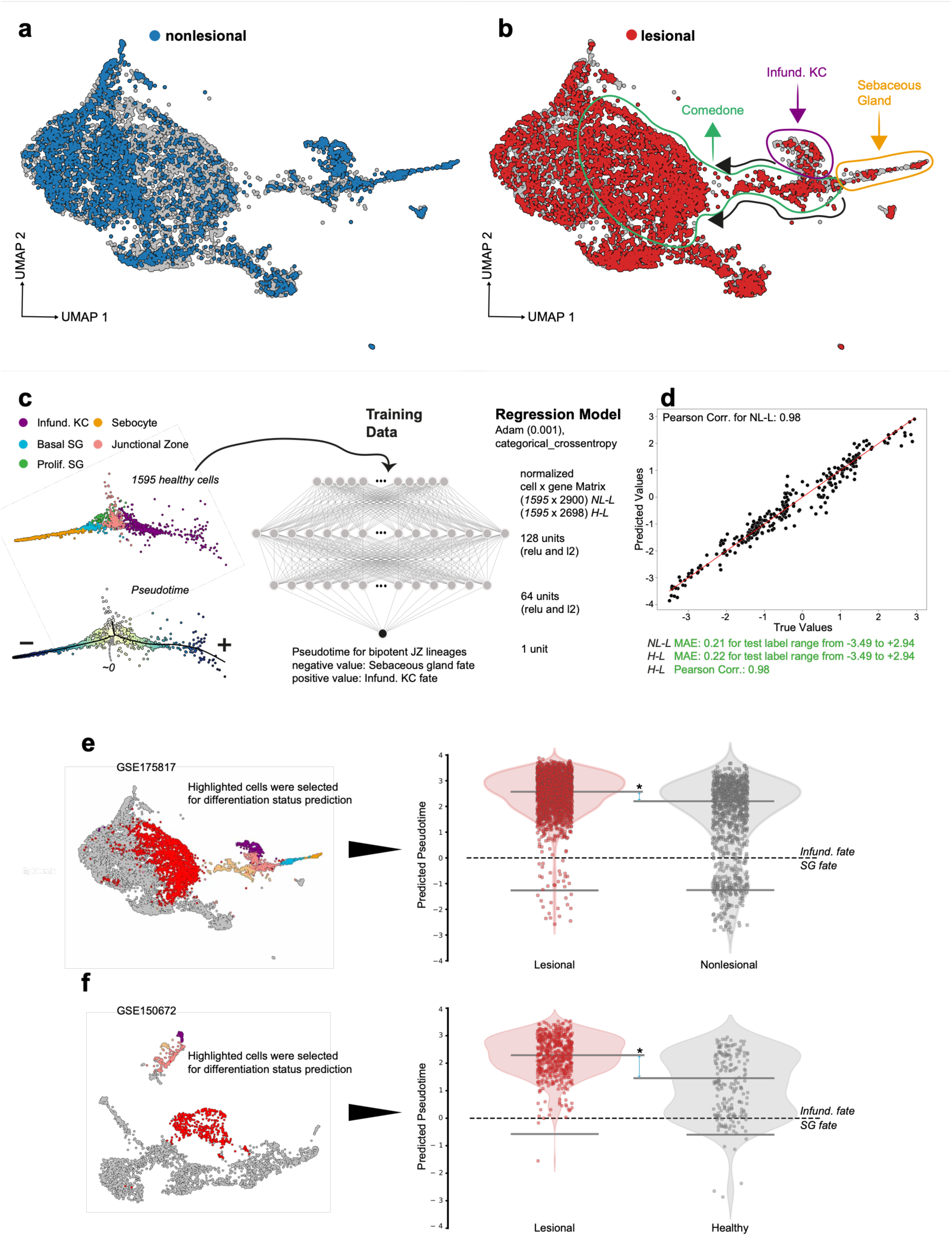
Pseudotime analysis supports a transcriptional shift toward infundibular-like differentiation in acne. **(a)** UMAP representation from Figure 3a of integrated acne samples highlighting cells derived from nonlesional skin. **(b)** Same embedding as in (a), highlighting cells derived from lesional acne skin. Lesional samples exhibited a marked reduction in sebocytes and infundibular keratinocytes, accompanied by expansion of the comedone population. **(c)** Schematic overview of the regression-based pseudotime framework used to model lineage trajectories originating from junctional zone cells. Opposing pseudotime directions were assigned to sebocyte and infundibular differentiation, enabling simultaneous inference of cell lineage identity and differentiation progression along both trajectories. **(d)** Performance evaluation of the regression model for predicting lineage-associated pseudotime values. **(e)** Predicted differentiation trajectories across cell populations in the integrated acne dataset. Comedonal keratinocytes exhibited enhanced progression along the infundibular differentiation trajectory, whereas sebocytes displayed attenuated differentiation. **(f)** Validation of lineage dynamics in an independent dataset comparing healthy and lesional acne skin. Similar shifts toward infundibular/comedonal differentiation and reduced sebocyte differentiation were observed. Abbreviations: H-L, healthy vs. lesional; JZ, junctional zone; KC, keratinocyte; NL-L, nonlesional vs. lesional; SG, sebaceous gland.

To test this hypothesis, we implemented a regression-based pseudotime model to infer lineage trajectories originating from junctional zone cells. Using the integrated embedding from (Düz et al., 2026b), we defined the junctional zone as the root population and mapped differentiation paths toward sebocytes and infundibular keratinocytes (Figure 5c). To distinguish the two lineages within a continuous pseudotime axis, we introduced a directional encoding by assigning opposite signs to the pseudotime values: positive values for the keratinocyte lineage and negative values for the sebocyte lineage. This enabled simultaneous modeling of both differentiation trajectories and cell fate decisions from junctional zone progenitors toward sebocytes or infundibular keratinocytes.

The model showed high performance in assessing cellular differentiation state (Figure 5d) and indicated that comedonal keratinocytes in lesional acne are shifted toward a more differentiated state (Figure 5e and f), consistent with excessive keratinization in the infundibulum, a well-established hallmark of acne development (Figure 1) (Vasam et al., 2023). Conversely, sebocytes in acne displayed attenuated differentiation, in line with a possible shift in lineage commitment. These trends were even more pronounced in another dataset comparing healthy and lesional acne samples (Figure 5f).

Collectively, our findings suggest that basal sebocyte progenitors in acne could adopt keratinocyte-like states, representing a potential mechanism underlying the initiation of comedone formation.

### Basal keratinocyte dysfunction and extended remodeling beyond the infundibulum in acne

To identify altered signaling pathways in lesion-specific keratinocyte clusters in acne and elucidate the interplay between the various cell populations at the nexus of comedogenesis, we performed a crosstalk analysis comparing different conditions. As aberrant keratinocytes occur exclusively in lesional skin, we compared these cells to normal infundibular keratinocytes. This comparison reflects the concept that comedone formation represents an abnormal differentiation trajectory of infundibular keratinocytes, consistent with our previous findings. Interestingly, across both comparisons (lesional acne vs nonlesional skin, and lesional acne vs healthy skin), pleiotrophin (PTN) signaling emerged as the most prominent difference, suggesting a role in early tissue remodeling during comedogenesis (Figure 6a and b). Consistent with previous reports implicating GATA6 in sebaceous duct homeostasis and acne pathogenesis (Oules et al., 2020), GATA6 expression was markedly reduced in lesional acne. In contrast, C10orf99 expression was restricted to lesional acne and absent from both healthy and nonlesional skin, identifying it as a disease-associated marker of comedonal keratinocytes (Figure 6b). Furthermore, reduced retinoic acid signaling was observed in the aberrant keratinocytes compared with infundibular keratinocytes in both datasets (Figure 6c). In healthy and nonlesional skin, basal keratinocytes of the IFE robustly expressed POSTN and ERRFI1 (Figure 6d and e). Strikingly, both genes were entirely absent from lesional skin.

**Figure 6.**
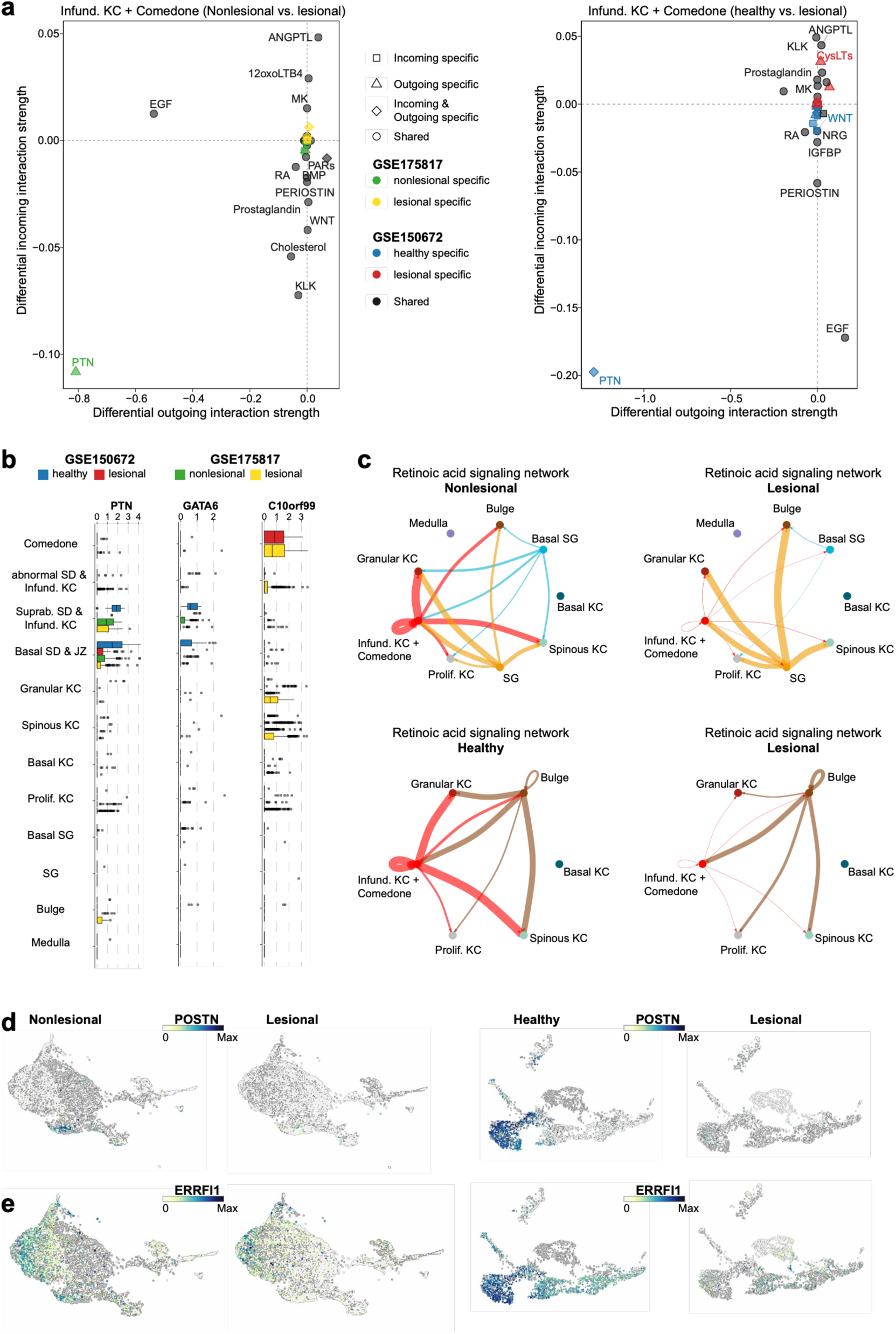
Altered signaling pathways and basal keratinocyte dysfunction in acne lesions. **(a)** Differential cell-cell communication analysis comparing lesional comedonal keratinocytes with infundibular keratinocytes. PTN signaling was consistently altered across independent acne datasets. **(b)** Differential expression of selected genes. Bar plots show expression changes of PTN, GATA6, and C10orf99 in lesional versus healthy skin (GSE150672) and lesional versus nonlesional skin (GSE175817). **(c)** Comparison of signaling pathway activity between lesional acne-associated cell states and their corresponding healthy or nonlesional counterparts, highlighting reduced retinoic acid signaling in comedonal cell states. **(d-e)** Expression of POSTN and ERRFI1 across healthy, nonlesional, and lesional skin cell populations. Both genes were expressed in basal interfollicular epidermal keratinocytes in healthy and nonlesional skin but were absent in lesional acne samples, indicating broader epithelial remodeling beyond the pilosebaceous unit. Abbreviations: JZ, junctional zone; KC, keratinocyte; SD, sebaceous duct; SG, sebaceous gland.

## 3 DISCUSSION

Microcomedone formation is considered one of the earliest detectable events in acne pathogenesis, yet the molecular “master switch” governing this process remains unresolved. Traditionally, factors such as comedogenic lipids, androgen, local cytokine imbalance, and bacterial colonization were implicated as causative agents (Contassot and French, 2014; Saurat, 2015; Shaheen and Gonzalez, 2013). However, the comedone switch hypothesis proposes that these factors converge on a cellular imbalance in the JZ, promoting a shift from sebaceous to infundibular differentiation (Clayton et al., 2019; Saurat, 2015; van Steensel, 2025; van Steensel and Goh, 2021).

While our observations are consistent with the comedone switch hypothesis, several limitations should be considered when interpreting these findings. The observed depletion of SG cells and expansion of comedonal keratinocytes were derived from scRNA-seq data, where cell capture is inherently stochastic and may be influenced by tissue dissociation, sampling variability, and differential cell recovery. Consequently, changes in cell proportions should not be interpreted as direct quantitative evidence of lineage conversion. Likewise, our analyses infer transcriptional relationships rather than direct cell fate transitions, as lineage tracing is not possible in human scRNA-seq datasets.

An additional limitation is that the analyzed datasets were generated independently, and the precise developmental stage of the sampled acne lesions is unknown. Lesional biopsies may contain varying proportions of comedones, papules, pustules, or mixed inflammatory lesions. Similarly, although the putative microcomedone-like state was identified in macroscopically nonlesional skin, its exact developmental stage cannot be determined. Thus, our findings should be interpreted as supporting a transcriptional model of early comedogenesis rather than defining a precise temporal sequence of lesion development.

Histological observations of atrophic SGs in lesional acne skin supported this hypothesis and associated it with LRIG1⁺ progenitor cells. A more recent study indicates that LRIG1 is not a specific marker of the human JZ, as it is broadly expressed across the PSU and IFE (Düz et al., 2026b). Using spatial transcriptomics, PTN was identified as a more specific human marker, expressed in basal cells of both the infundibulum and the JZ (Düz et al., 2026a). Notably, altered PTN signaling emerged as the most prominent difference between abnormal keratinocytes and neighboring infundibular and isthmus cells (Figure 6a), suggesting a potential role of this pathway in the comedone switch. Interpretation of this finding should, however, be made with caution. The lesional keratinocyte cluster likely comprises not only comedonal keratinocytes but also neighboring interfollicular epidermal cells, reflecting the difficulty of separating these closely related epithelial populations based solely on transcriptomic profiles. Consequently, differences in PTN signaling may partially reflect changes in cellular composition rather than comedone-specific biology.

The reprogramming observed in comedones is characterized by increased expression of keratinization-associated genes such as KRT6 (Cunliffe et al., 2000) and a selective loss of differentiation markers like KRT79 in the infundibular compartment (Fontao et al., 2020), while KRT17 expression remains preserved (Supplementary Figure S5a-c) (Veniaminova et al., 2013). This altered expression pattern coincides with markedly reduced levels of the transcription factor GATA6, a key regulator of JZ homeostasis and sebaceous lineage specification, suggesting that loss of GATA6-mediated control underlies the observed transcriptional shift (Oules et al., 2020). Consistent with this, mouse models lacking GATA6 develop cystic infundibular lesions (Swanson et al., 2019). Retinoid signaling represents an additional layer of control. Retinoids upregulate GATA6, thereby likely restoring ductal differentiation capacity (Oules et al., 2020). Our data further suggest a local vitamin A deficiency in the aberrant keratinocytes (Figure 6c).

Given the remodeling nature of the upper hair follicle during comedogenesis, we next asked how far comedonal alterations might extend within the tissue. The infundibulum is an invaginated structure of the upper follicle, yet comedones are macroscopically visible on the skin surface. This suggests that pathological signaling originating from the infundibulum may extend beyond the follicular boundary, likely affecting neighboring IFE in a gradient-like manner. We hypothesize that comedonal keratinocytes within the infundibulum and adjacent IFE adopt similar transcriptional programs.

This potential relationship prompted us to investigate whether basal alterations might also occur within the IFE. Indeed, in healthy and nonlesional skin, basal keratinocytes of the IFE robustly expressed POSTN and ERRFI1 (Figure 6d and e). Strikingly, both genes were entirely absent from lesional skin, indicating a shared basal vulnerability in acne that extends beyond the infundibulum. As basal keratinocytes in nonlesional skin retain expression elsewhere, we infer that the loss of these markers is likely spatially restricted to IFE regions immediately surrounding the infundibulum. Given that ERRFI1 encodes an inhibitor of EGFR signaling (Cairns et al., 2018; Frosi et al., 2010; Reschke et al., 2010), its absence may contribute to enhanced EGF pathway activity in acne-prone regions, which has previously been linked to increased sebum production (Dahlhoff et al., 2014). Supporting this, genome-wide association studies have identified ERRFI1 variants among acne risk loci (Mitchell et al., 2022; Teder-Laving et al., 2024), underscoring the genetic contribution to basal pathway dysregulation.

Environmental factors, including activation of the aryl hydrocarbon receptor (AhR) pathway, may also contribute. AhR activation can suppress OVOL1, a transcription factor that normally inhibits canonical Wnt signaling and promotes epidermal differentiation (Schneider et al., 2014). Loss of OVOL1 expression in comedonal cells may thus relieve this inhibitory brake (Supplementary Figure S5d), amplifying hyperproliferative and keratinizing programs. Notably, OVOL1 is also implicated as an acne risk locus in GWAS (Petridis et al., 2018; Teder-Laving et al., 2024).

Together, microbial and environmental triggers converge on shared signaling pathways, potentially disrupting the balance between differentiation and proliferation within the upper follicle. However, the temporal relationship between these processes remains unresolved, and it is unclear whether inflammatory signaling primarily precedes, accompanies, or reinforces the lineage imbalance associated with comedogenesis. Although only a minority of follicles develop visible lesions, histological analyses reveal that microcomedones are already present in a substantial fraction of clinically normal PSUs (Cunliffe et al., 2000; Fontao et al., 2020; Saurat, 2015). The skin’s ability to resolve most comedones within days highlights its regenerative potential, emphasizing that therapeutic strategies should focus on preventing naïve PSUs from undergoing the comedone switch.

Retinoids remain the most effective intervention (Layton, 2009), yet their systemic and broad epithelial effects call for more selective strategies particularly to reduce side effects (Draghici et al., 2021; McLane, 2001). Targeted modulation of the *C. acnes* niche, controlled reduction of sebum production, and restoration of local retinoid and EGFR signaling balance may act synergistically to stabilize PSU differentiation. While our findings provide transcriptomic support for the comedone switch hypothesis and identify candidate pathways associated with this process, future longitudinal, spatially resolved, and functional studies will be required to establish causal mechanisms and the temporal sequence of events during human comedogenesis. Such studies may ultimately facilitate the development of more targeted preventive and therapeutic strategies for acne.

## 4 MATERIALS AND METHODS

### Immunohistochemistry

Paired 4-mm punch biopsies of lesional and nonlesional skin were obtained from the left upper back of a patient with clinically diagnosed acne vulgaris (BioIVT, Westbury, NY, USA). Healthy donor skin biopsy was kindly provided by the Institute of Allergology, Charité – Universitätsmedizin Berlin (Berlin, Germany). All samples were formalin-fixed, paraffin-embedded (FFPE) and sectioned at 4 μm.

A standardized immunohistochemistry protocol was followed. Following deparaffinization and rehydration, heat-induced epitope retrieval was performed in pH6.0 buffer (Dako; Cat. No. S2369) at 95°C for 15 min. Endogenous peroxidase activity was blocked with 0.3% hydrogen peroxide, followed by protein blocking (Dako; Cat. No. X0909). Skin sections were incubated overnight at 4°C with primary antibodies against human-S100A7A (Invitrogen, Cat. No. PA5-104043, RRID: AB_2853373; 1:1000) and human-SERPINB4 (Invitrogen, Cat. No. MA5-25407, RRID: AB_2725329; 1:200). Detection was performed using EnVision+ HRP-labelled polymer secondary antibodies (Dako; Cat. No. K4003 and K4001, respectively) and the ImmPACT AEC substrate kit (Vector Laboratories; Cat. No. SK-4205). Sections were counterstained with Gill’s hematoxylin I and mounted using an aqueous mounting medium.

Images were acquired at 20× magnification using a Keyence BZ-X810 microscope system (Keyence Corporation, Osaka, Japan) and analyzed using BZ-X Analyzer software.

The use of healthy control tissue was approved by the Ethics Committee of Charité – Universitätsmedizin Berlin (approval number EA/064/23). Commercially sourced acne tissue specimens were obtained from BioIVT in accordance with FDA guidelines.

### Data Processing and Integration of Acne scRNA-seq Datasets

The GSE175817 dataset (NL–L) is derived from punch biopsies collected from the backs of six acne patients. For each individual, samples were obtained from lesional skin (directly from an acne lesion) and patient-matched, adjacent nonlesional skin. Nonlesional skin refers to tissue that appears macroscopically healthy but is taken from skin directly surrounding the lesion. The GSE150672 dataset (H–L) comprises lesional samples from acne patients and site-matched healthy control samples from individuals without acne.

For GSE175817, we used the authors’ processed Seurat object, which includes a coarse annotation of major cell types. From this object, we isolated the epidermal cluster and performed subclustering using our analysis pipeline to obtain a more fine-grained classification of epidermal cell states.

Preprocessing was performed in Seurat using default parameters unless otherwise specified. A lenient filtering strategy was applied, retaining cells with fewer than 25% mitochondrial reads and more than 300 detected genes. The data were log-normalized, and highly variable features were identified. Prior to dimensionality reduction, the data were scaled, and mitochondrial gene content was regressed out. Principal component analysis (PCA) was then performed, and the nearest-neighbor graph was constructed using the first 20 principal components. Clustering was carried out using the Louvain algorithm, and the resulting structure was visualized using UMAP.

Since substantial batch effects between conditions were observed, batch correction was performed using Harmony, specifying condition and donor as covariates. Neighbors, clusters, and UMAP embeddings were subsequently recalculated. The initial lenient filtering and broad subsetting resulted in the emergence of low-quality clusters, which were identified based on differential gene expression (DGE) analysis as well as characteristic quality metrics, including low numbers of detected features and elevated mitochondrial read fractions. These clusters were removed, and the full workflow (scaling, integration, clustering, and UMAP) was rerun, resulting in a final Seurat object that clearly resolved epidermal and PSU cell types. This analysis also revealed a previously undescribed keratinocyte subpopulation with a distinct transcriptomic signature. All remaining cell types were annotated using marker genes reported in the literature. For the embedding restricted to nonlesional cells, the same analytical strategy and quality-control criteria were applied.

For the GSE150672 dataset, the processed count matrix was downloaded, filtered for acne versus healthy conditions, and processed analogously to the pipeline described above.

### Nonlesional microcomedone and healthy Classifier

To train a classifier for predicting microcomedone-related cell states in the two embeddings from Figure 3a (GSE175817 and GSE150672), we imported the *h5ad* object corresponding to the nonlesional embedding into a Jupyter Notebook. For each dataset, the corresponding embedding was loaded and subset to the set of highly variable genes shared with the nonlesional reference embedding, and an independent neural network classifier was trained. Gene expression matrices were extracted as dense arrays and cell-type labels were encoded using a label encoder, followed by one-hot transformation. The data were split into training, validation, and test sets using an 80/20 split for test data and an additional 80/20 split within the training portion for validation, stratified by the encoded labels.

A fully connected neural network was implemented using Keras. The model consisted of two hidden layers (128 and 64 units) with *ReLU* activation and L2 regularization (λ = 0.001), followed by a softmax output layer. The network was trained using the *Adam* optimizer (learning rate = 0.001) and categorical cross-entropy loss. Early stopping (patience = 10 epochs) and model checkpointing based on validation loss were applied during training. Models were trained for up to 300 epochs with a batch size of 64 using multiprocessing.

The optimal number of epochs was determined by identifying the minimum validation loss. Subsequently, the model was retrained on the full training set using 120% of the initially determined optimal epoch count. Performance was evaluated on the held-out test set using macro-averaged precision, recall, and F1-score. The final models were applied to the two embeddings to obtain predicted cell-type labels and to assess whether lesional cell states from acne datasets exhibit transcriptional similarity to the putative microcomedone-like transcriptional state.

For the healthy classifier, the embedding from study (Düz et al., 2026b) was used, and the same preprocessing steps, model architecture, and training procedure described above were applied.

### Mapping to the HSCA Core Using scArches

We mapped all query datasets onto the HSCA core reference using scArches (v0.6.1), following the published workflow for reference mapping (https://docs.scarches.org/en/latest/hlca_map_classify.html). This approach enables a joint latent embedding of reference and query datasets and provides a stable basis for downstream analyses, including clustering and label transfer.

The query datasets comprised three conditions (acne, nonlesional, and healthy), which were mapped independently. For each condition, only genes used as input features in the HSCA reference model were retained; genes absent in the reference were padded with zeros as required by the scArches framework.

Reference mapping was performed using scVI with the surgery procedure (Lopez et al., 2018). During surgery, sample-specific batch adapters were inserted into the frozen HSCA reference model, and only these adapter parameters were updated. For each query condition, adapter training was run for up to 500 epochs with early stopping enabled and weight decay set to 0. All models were saved for reproducibility.

### Label Transfer and Uncertainty Estimation

Cell-type annotations from the HSCA core were propagated to the mapped query datasets using the k-nearest neighbour (kNN) label transfer routine implemented in scArches (Lotfollahi et al., 2022). The latent representations of HSCA reference and query cells were concatenated to construct a joint embedding, from which a kNN graph (k = 50) was built. Each query cell received the most frequent HSCA reference label among its neighbours, and an uncertainty score was computed as described previously.

All uncertainty scores were stored and linked to the corresponding cells in the embeddings displayed in Figure 3.

### Pseudotime Modeling of Junctional Zone Differentiation

To infer lineage trajectories originating from junctional zone progenitors, we implemented a regression-based pseudotime model. Using the integrated embedding from Düz et al. (2026b), the JZ was defined as the root population, and differentiation paths toward sebocytes and infundibular keratinocytes were mapped. To model both lineages along a continuous pseudotime axis, we assigned opposite signs to the pseudotime values: positive for the keratinocyte lineage and negative for the sebocyte lineage. This encoding allowed simultaneous modeling of both differentiation trajectories and cell fate decisions from JZ progenitors.

For each dataset, embeddings corresponding to infundibular, comedonal, JZ, and SG lineages were subset to the highly variable genes shared with the healthy reference embedding. Independent neural network regressors were trained on the embedding. Gene expression matrices were extracted as dense arrays, and pseudotime values were used as regression targets. Data were split into training, validation, and test sets (80/20 split for test data, and an additional 80/20 split of the training data for validation).

A fully connected neural network was constructed using Keras, consisting of two hidden layers (128 and 64 units) with ReLU activation and L2 regularization (λ = 0.001), and a linear output layer for pseudotime prediction. The network was trained with the Adam optimizer (learning rate = 0.001) using mean squared error as the loss function. Early stopping (patience = 30 epochs) and model checkpointing based on validation loss were applied. Models were trained for up to 500 epochs with a batch size of 64 and multiprocessing enabled. The optimal number of epochs was determined based on minimum validation loss, and the model was subsequently retrained on the full training set using 120% of the optimal epoch count.

Model performance was evaluated on held-out test sets using mean absolute error and Pearson correlation between predicted and true pseudotime values. Final pseudotime predictions were added to the metadata of the corresponding cells. Lesional, nonlesional, and healthy cells were separated, and target clusters were defined for lineage-specific analyses. Predicted pseudotime values were visualized using combined strip and violin plots, with median pseudotime values indicated for positive and negative lineages. We used a Mann-Whitney-U test to compare the pseudotime distributions of lesional versus nonlesional/healthy cells, separately for positive and negative pseudotime values. These visualizations facilitated comparison of pseudotime distributions between acne lesional and nonlesional or healthy cells, enabling assessment of differentiation biases toward sebocytes or infundibular keratinocytes.

### Cell-Cell Crosstalk Analysis

To investigate altered signaling pathways in lesion-specific keratinocyte clusters in acne, we performed a crosstalk analysis comparing different conditions. As aberrant keratinocytes are exclusive to lesional skin, these cells were compared to normal infundibular keratinocytes to assess differential signaling associated with comedone formation, which is considered an abnormal differentiation trajectory of infundibular keratinocytes. For this purpose, comedone cells and abnormal SD/JZ keratinocytes were technically relabeled as “Infundibular keratinocytes” in the metadata, allowing the analysis to contrast normal versus lesional keratinocytes within the same reference cell type.

Cell-cell communication analysis was performed using CellChat following the standard workflow with default parameters as described by Jin et al. (2025). We used the extended CellChat database (version 2), excluding extracellular matrix–receptor interactions and direct cell-cell contact pathways, as these interactions (e.g., collagen, laminin, tenascin) were overwhelmingly abundant in skin samples and could obscure other signaling pathways.

The computeCommunProb function was used to calculate communication probabilities, applying a truncated mean with a trimming parameter of 0.10 to avoid overly stringent filtering. During downstream analysis with netAnalysis_signalingChanges_scatter(), CypA signaling was excluded.

This approach allowed the identification of signaling pathways specifically altered in lesion-associated, infundibular-like keratinocytes compared with normal infundibular keratinocytes.

## DATA AND CODE AVAILABILITY

All analyses were performed using publicly available datasets (GSE175817 (Do et al., 2022) and GSE150672 (Hughes et al., 2020)), software and packages. All scripts are available upon request from the corresponding author, TD.

## CONFLICT OF INTEREST

TD, TR, HR, BA, SG, and NH are employees of Beiersdorf AG. DT and JB received consultation fees from Beiersdorf AG. The remaining authors state no conflict of interest.

## ACKNOWLEDGEMENTS

This work was funded by Beiersdorf AG. We thank Dr. Stefan Frischbutter (Institute of Allergology, Charité – Universitätsmedizin Berlin) for facilitating access to healthy skin tissue samples used in this study.

## AUTHOR CONTRIBUTIONS

Conceptualization: TD, NH; Formal Analysis: TD; Funding Acquisition: SG, BA, HR; Resources: TD; Software: TD; Supervision: NH, JB, DT; Visualization: TD; Writing – Original Draft Preparation: TD. Antibody staining experiments: TR.

## DECLARATION OF ARTIFICIAL INTELLIGENCE/LARGE LANGUAGE MODEL USE

During the preparation of this work, the author(s) used ChatGPT in order to assist with language refinement, phrasing, and general text formulation. The tool was primarily used to improve clarity, grammar, and readability of the manuscript. After using this tool, the author(s) critically reviewed and edited all generated content as needed and take full responsibility for the final content of the publication.

---

**Supplementary Figure S1.**
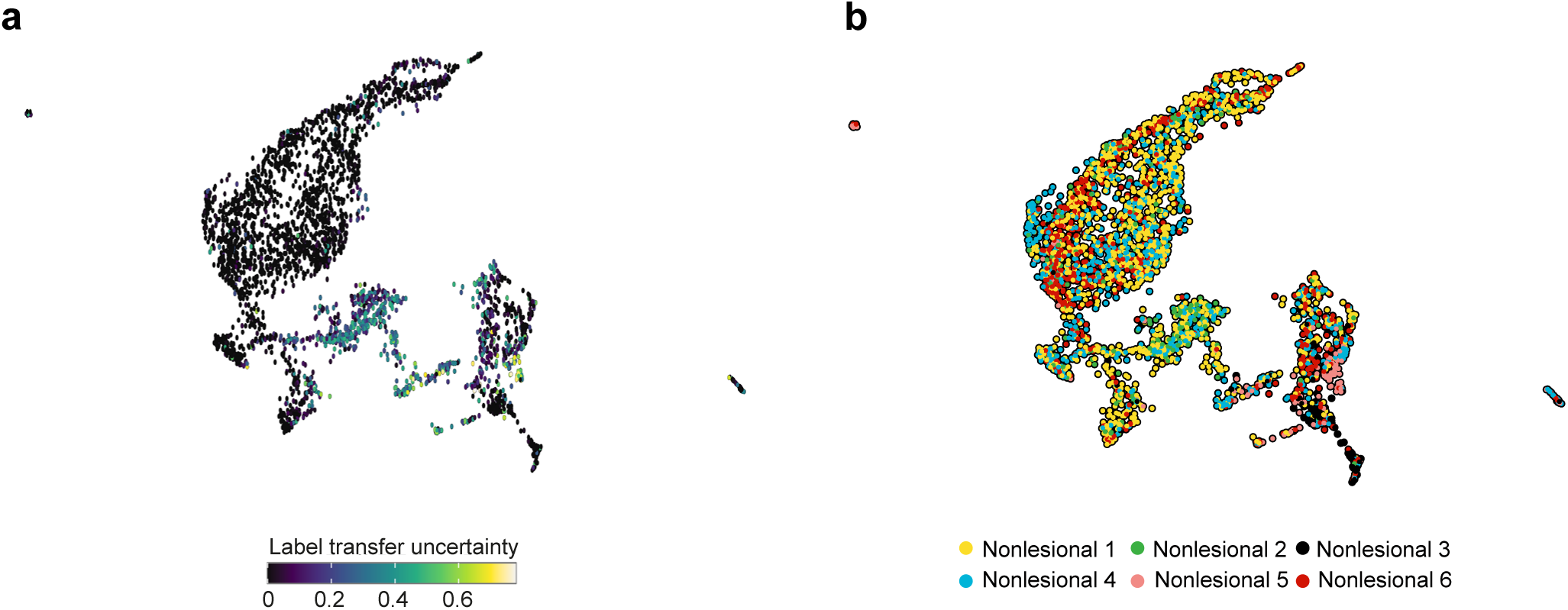
: The putative microcomedone population is absent from healthy skin and enriched in a subset of acne patients. (a) Projection of the nonlesional acne skin dataset onto the healthy core Human Skin Cell Atlas (∼300,000 cells). The putative microcomedone cluster identified in nonlesional acne skin exhibited high cell type label transfer uncertainty when projected onto the healthy reference atlas, suggesting that it represents a cell state absent from healthy skin. (b) Distribution of cells belonging to the putative microcomedone cluster across individual patients. The population was predominantly contributed by three patients, indicating inter-individual variability in the abundance of this cell state.

**Supplementary Figure S2.**
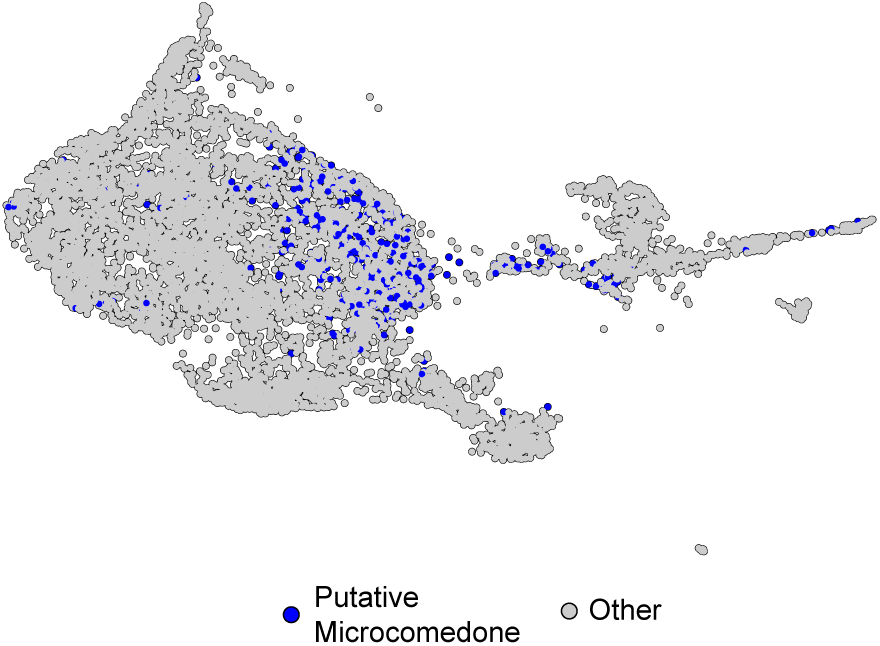
: The putative microcomedone population identified in Figure 2c localizes to a lesional acne-associated region of the integrated embedding.

**Supplementary Figure S3.**
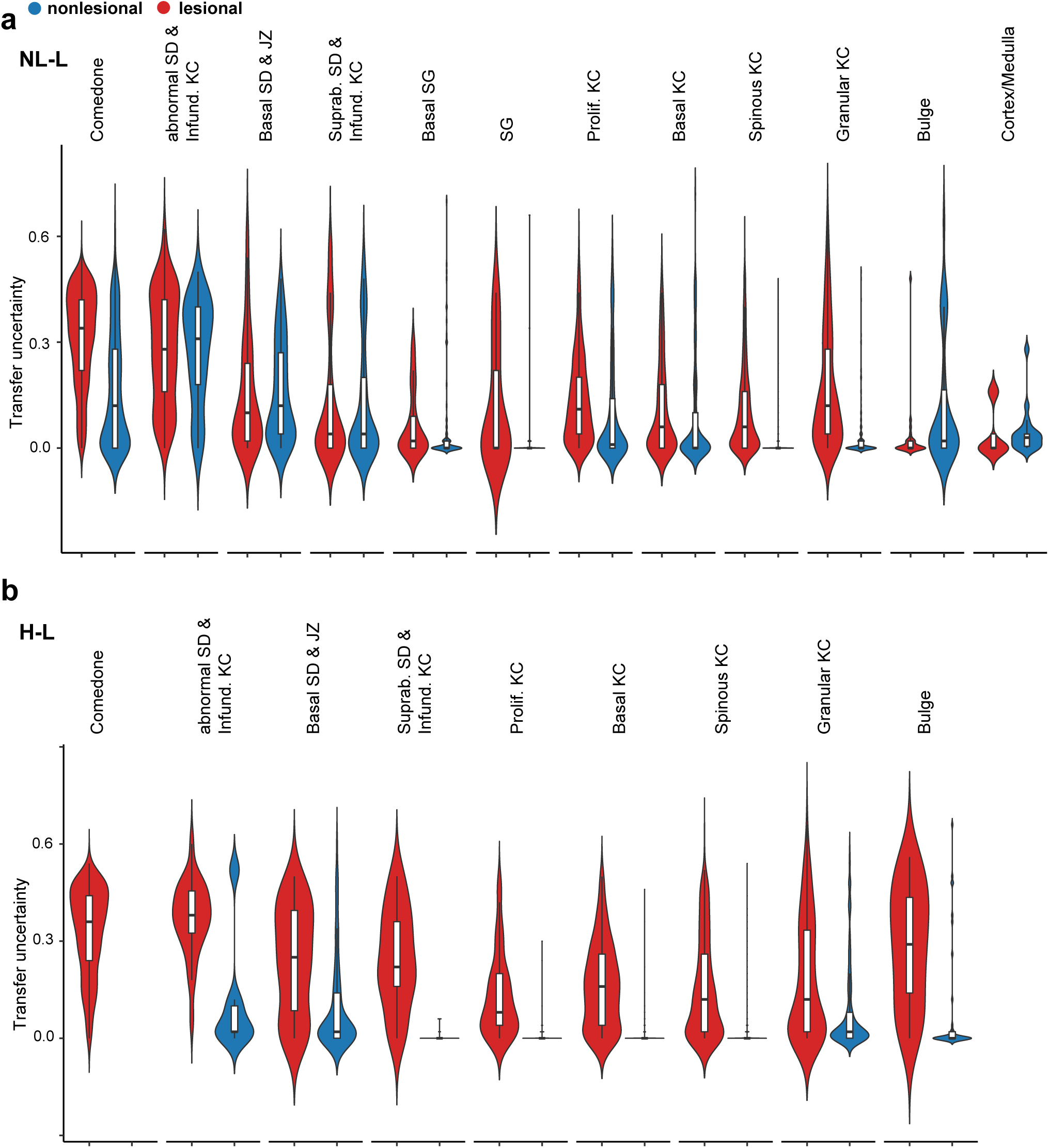
: Transfer learning uncertainty across skin cell populations. Violin plots showing transfer-learning uncertainty scores for selected skin cell populations following projection onto the healthy HSCA reference. Higher uncertainty indicates reduced correspondence to cell states present in healthy skin. Putative comedonal keratinocytes displayed the highest uncertainty scores, whereas established epidermal, sebaceous gland, and hair follicle populations mapped with substantially lower uncertainty, supporting the presence of an acne-specific cell state not represented in healthy skin. Abbreviations: H-L, healthy vs. lesional; JZ, junctional zone; KC, keratinocyte; NL-L, nonlesional vs. lesional; SD, sebaceous duct; SG, sebaceous gland.

**Supplementary Figure S4:**
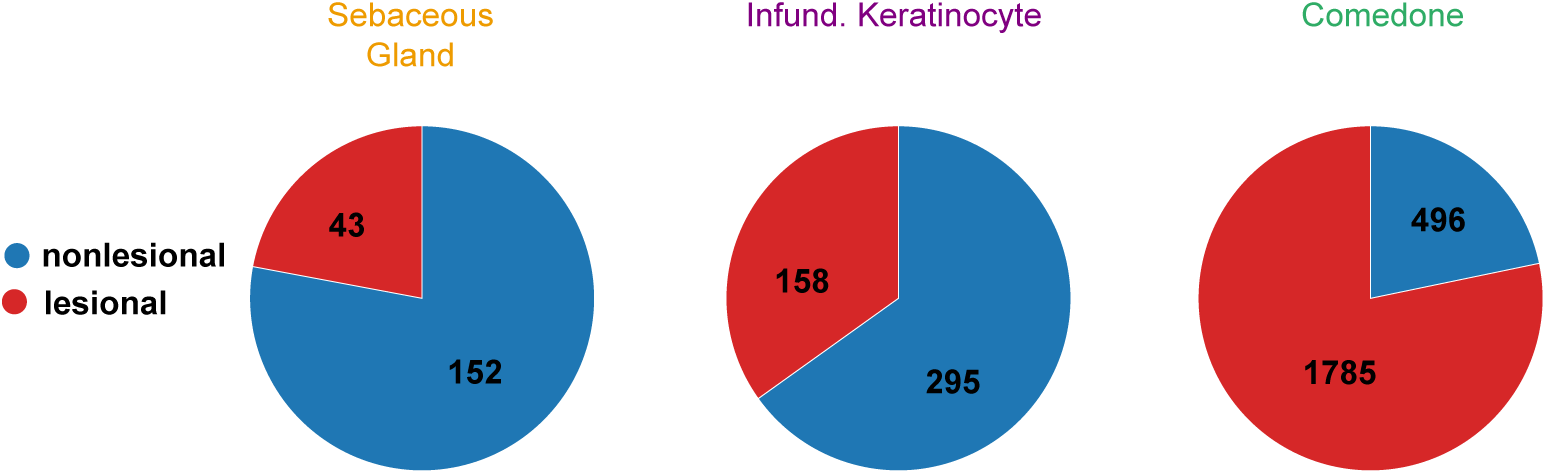
Quantification of cell populations associated with comedogenesis. Pie charts showing the distribution of sebaceous gland cells, infundibular keratinocytes, and comedonal keratinocytes in lesional and nonlesional skin. Lesional samples exhibited a marked reduction in sebaceous gland cells and infundibular keratinocytes, accompanied by a substantial expansion of comedonal keratinocytes. Notably, the comedonal keratinocytes detected in nonlesional skin corresponded to the putative microcomedone cluster identified in our previous analyses (Supplementary Figure S2).

**Supplementary Figure S5:**
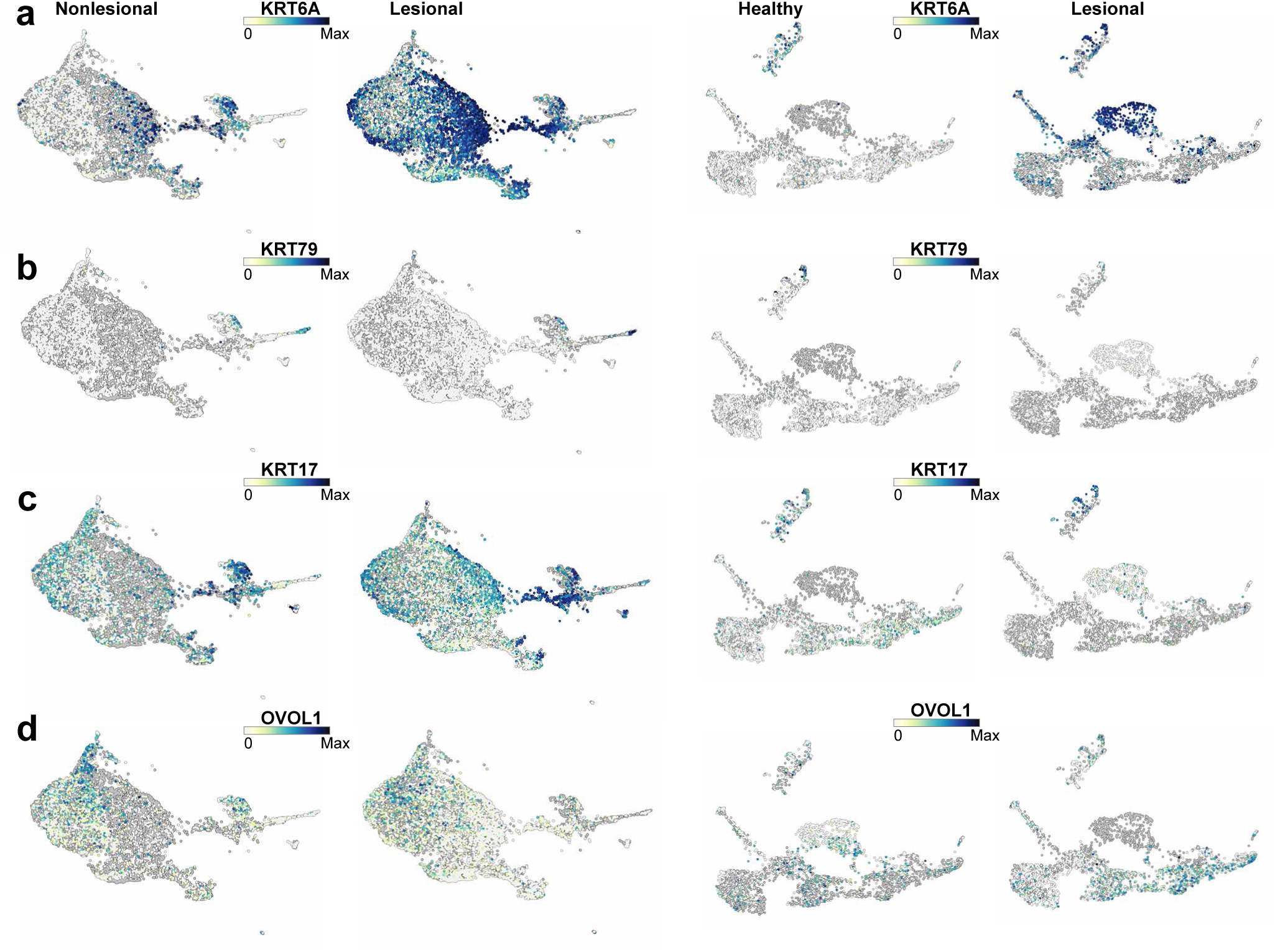
Expression of KRT6A, KRT79, KRT17, and OVOL1 in healthy, nonlesional, and lesional skin. **(a)** KRT6A expression. **(b)**, KRT79 expression. **(c)**, KRT17 expression. **(d)**, OVOL1 expression. KRT6A expression was increased in lesional comedonal populations, whereas KRT79 and OVOL1 expression were reduced. In contrast, KRT17 expression remained preserved. These patterns are consistent with the transcriptional reprogramming of comedonal keratinocytes associated with enhanced keratinization and altered differentiation.

## Notes

### Summary of Updates

This version of the manuscript has been revised to include additional antibody staining experiments validating markers identified in the comedone signature. The manuscript text has also been refined for clarity, and minor errors, including reference and formatting issues, have been corrected.

